# Dynamic Disorder in Chlorophyll Aggregation and Light-Harvesting Complex II in the Plant Thylakoid Membranes using Coarse-Grained Simulations

**DOI:** 10.1101/2025.02.18.638782

**Authors:** Renu Saini, Avinash Garg, Ananya Debnath

## Abstract

The dynamics of the aggregated light-harvesting complex (LHCII) associated with its antennae pigments can be crucial for a transition between light harvesting and dissipative states pivotal for non-photochemical quenching (NPQ). To this end, aggregation of chlorophyll-a (CLA) without the LHCII and pigment binding LHCII monomers in the plant thylakoid membranes have been investigated using coarse-grained molecular dynamics simulations at 293 K. Both CLA without the LHCII and pigment-binding LHCII monomers dynamically form and break dimers and higher-order aggregates in thylakoids within the simulation time. The contact lifetime and waiting time distributions of CLA dimers exhibit multiple time scales including most populated fast time scales and less populated slow time scales. The survival probability of CLA dimer in the absence of the LHCII follows a non-exponential decay with multiple residence time scales, leading to a time-dependent rate, unlike conventional rate theory. Such non-exponential decay of survival manifests the emergence of dynamic disorder in CLA without the LHCII resulting from the coupling between time scales of dimer formation and higher-order aggregates. The conformational fluctuations of the LHCII known for inter-CLA coupling variation occur on multiple time scales comparable to the LHCII dimer residence time scales leading to less probable but comparable and more probable slower inter-CLA fluctuations. This indicates the dynamic coupling in the LHCII conformations and their aggregates with the antennae pigments can result in dynamic disorder which will be highly relevant for the light-harvesting efficiency and regulation of NPQ.

**TOC Graphic:** 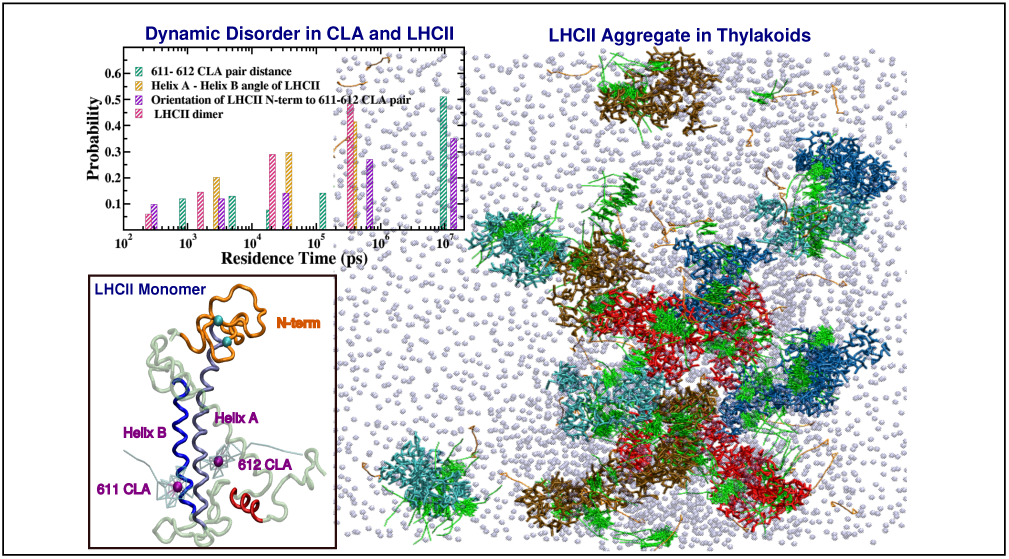

## Introduction

CLA is an essential pigment for photosynthesis, the mechanism through which plants transform light energy into chemical energy, fueling life. CLA pigments are associated with the light-harvesting complex II (LHCII), a protein trimer in photosystem II, through specific binding sites.^1,2^ The integrity of the trimeric conformation of LHCII is maintained via salt bridges and synergistic hydrogen bonds between the trimerization motif (WYR) and the pigments or lipids through the intermediate residues within the monomer. ^3^ Aggregates of LHCII facilitate the formation of CLA-CLA homodimers or CLA-lutein heterodimers, act as the quenching centers within LHCII, and are essential for photoprotection.^4^ The aggregation causes the dispersion of excitation energy as heat and participates in NPQ. ^5^ Aggregates of LHCII play an important role in maintaining the dissipative state of LHCII under high light conditions by redirecting the excess energy to the violaxanthin cycle ^5–7^ to prevent plants from photodamage.^8^ The primary site of light-dependent reactions of CLA binding LHCII is the thylakoid membranes of plant chloroplasts. ^9^ Plant thylakoids contain a mixture of unsaturated lipids distinguished by the number and location of the double bonds in their fatty acid chains. The critical ratio of the lipids with less and a higher number of unsaturation in thylakoid provides stability to the CLA dimer. ^10^ Experimental evidence from fluorescence spectroscopy at 77 K shows that lipids in thylakoid membrane regulate LHCII organization and that LHCII aggregation stabilization is mediated by selective lipids (MGDG and DGDG).^4^

CLA aggregates and their derivatives have therapeutic applications in photodynamic therapy (PDT) when bound to membranes without the LHCII.^11–14^ CLA can induce detoxification, aid in the healing of wounds, and contribute to overall wellness.^15–17^ Several experimental techniques, such as fluorescence and circular dichroism, nuclear magnetic resonance self-diffusion, transmission electron microscopy, and adsorption-stripping cyclic voltammetry, are used to investigate the spectral properties of CLA aggregates and suggest that CLA forms aggregate structures like micelles in pure water solvent due to the hydrophobic effect caused by its phytol tail by minimizing the repulsive interactions between water and phytol tail.^18^ Fluorescence spectroscopy study suggests that the self-assembly of CLA is facilitated by hydrogen-bonding ability and dipole-dipole interactions of the solvent–water mixtures.^19^ The solvent ligands attached to the Mg atoms are substituted by water, allowing the formation of aggregates.^20^ An all-atom molecular dynamic simulation was performed to investigate the structural and dynamic characteristics of chlorophyll in several organic solvents, including water, methanol, and benzene. The oxygen atom of solvent molecules forms a bond with the central Mg atom of the chlorin ring.^21^ Further, our coarse-grained (CG) molecular dynamics (MD) study supports the fact that CLA forms higher order aggregates in DPPC and plant thylakoid membranes, where van der Waals interactions facilitate the aggregate formation.^22,23^

A two-dimensional electronic spectroscopy study reveals that decay in frequency fluctuation correlation function (FFCF) for CLA in three distinct solvents, tetrahydrofuran, diethyl ether, and methyl alcohol, follow a multiexponential function using the central line slope method. The CLA molecule exhibits the fastest relaxation time, ranging from 0.2 ps to 0.7 ps, and the slowest time scale is greater than 1 ns in all solvents.^24^ All-atom MD simulations demonstrate that the residence time of water molecules in the chlorin ring follows the multi-exponential distribution with three characteristic time scales 10, 200, and 1300 ps.^21^ Single-molecule spectroscopy study uncovers the presence of dynamic equilibrium between light-harvesting and dissipative conformations.^25^ The potential sites responsible for conformational switching in LHCII of Chlamydomonas reinhardtii are identified by using NMR spectroscopy.^26^

The antennae pigments surround the light-harvesting protein in a belt-like arrangement with specific positions and orientations. The protein-pigment interactions and pigment-pigment interactions change the light-harvesting properties. The protein serves as a highly heterogeneous dynamic environment to the pigment bound at a specific site of the protein. Any dynamical change in the protein occurring at a wide distribution of time scales affects the transition energy of the pigment, which in turn, changes the inter-pigment coupling. The rapid, collective nuclear vibrational modes of the protein and pigments are referred to as dynamic disorder,^27^ interact with the electronic excited states of the pigments. On the other hand, the slower, larger-amplitude slow structural fluctuations of proteins, referred to as static disorder, can significantly impact the light-harvesting efficiency of the pigment-protein complex or LHCII.^27,28^ Structural fluctuations or disorders in proteins may enable LHCII to switch the antennae between an efficient energy dissipation state and an effective light-harvesting state.^8,27,28^ This intrinsic disorder lets the protein access various substrates in its energy landscape, crucial for efficient energy regulation in photosynthesis through NPQ.^29^ NPQ of chlorophyll fluorescence regulates the function of the light-harvesting antenna of photosystem II (PSII).^30^ NPQ is primarily activated under excess illumination, the conformation of LHCII switches from the light-harvesting to the quenched state that dissipates excess energy as heat, protecting the photosynthetic equipment. Moreover, under excess illumination, the LHCII monomer and trimer show aggregation behavior.^31,32^ These aggregations can be helpful in the xanthophyll cycle by changing the interaction lipids to LHCII. A previous study^29^ demonstrates that LHCII exhibits a dynamic function that relies not on significant conformational transitions between distinct states, but rather on alterations in pigment configurations and/or environmental variations stemming from the intrinsic disorder of these complexes. Single-molecule spectroscopy study uncovers the presence of dynamic equilibrium between light-harvesting and dissipative conformations.^25^ Intrachromophore dephasing arises from static and dynamic disorder, driven by the slow and fast conformational dynamics, respectively of the aggregates of the photosynthetic light-harvesting complexes.^33^ Dynamic disorder due to the transitions between different excitons of the chromophore in LHCII causes intra-chromophore dephasing. ^33^ Dynamic or static disorder, important in LHCII, is common in proteins and may play significant roles in other biological processes, including enzymatic reactions.^34–36^

Although LHCII conformational fluctuations and their coupling to phonons are important in energy harvesting and dissipation, the conformational transitions of the protein pigment systems between these different states are not understood yet. Although LHCII aggregates indicate quenched states,^30^ a single LHCII crystal also confirms the existence of the active quenched state.^37^ For this, no major conformational switch is anticipated in the LHCII. Therefore, the chlorophyll distance distribution can be one of the crucial properties to get more insights into the quenched state. On the other hand, the structure of the LHCII in the presence of a membrane differs significantly from the crystal structure and is known to be associated with the experimental light-harvesting state.^37^ Molecular dynamics simulations show that the N-terminus of the LHCII in the membrane is highly disordered and strongly correlated with the energetic disorder of the lowest energy site of the complex, Chla611-Chla612. Under stressed conditions, lipids in the membrane can reorganize impacting the N-terminal fluctuations of the LHCII which can cause a large variation in Chla611-Chla612 coupling.^37^ Earlier simulations demonstrate different conformational states of the LHCII characterized by significant variations in the coupling strength between pigments. This strongly supports the hypothesis that the NPQ transitions can be governed by the conformational variations of LHC complexes through pigment coupling. Since distances between pigments are the most sensitive and crucial indicators of conformational flexibility of the LHCII relevant for the light-harvesting or energy-dissipating states, the current study, therefore, focuses on the detailed dynamic aspects of chlorophyll aggregation that lays the foundation to investigate the principles of dynamic disorder in a more complex system of LHCII. It is not clear how pigment-pigment distance fluctuations are related to the conformational fluctuations of LHCII and their relation to the aggregation behavior. The contribution of dynamic disorder in CLA or LHCII aggregation remains unclear and the role of dynamic disorder in light harvesting and excess energy dissipation remains unresolved. Thus, the current study aims to delve into the intricate dynamics of CLA aggregation in plant thylakoid membranes using M3^22^ and Martini model^38^ of CLA. The study of CLA aggregation builds a foundational understanding of the dynamic disorder in a simpler system where CLA molecules dynamically form dimers and higher-order aggregates and break and reform. The formation and breakage mechanism of a dimer can affect the formation and breakage mechanism of a trimer which can again affect the same for the higher-order aggregates. Thus CLA dimer dynamics is a more complex dynamic process than that for the usual two-state systems and can be coupled to the dynamics of multi-state higher-order aggregates. The contact lifetime and waiting time distribution of CLA dimers follow a broad range of time scales. The decay in survival probability and rate of CLA dimer exhibits a non-exponential behavior suggesting the presence of dynamic disorder in CLA aggregation. LHCII monomers with pigments embedded in the thylakoid membrane dynamically form dimers and higher-order aggregates. The multi-exponential decay in survival probability of the LHCII dimer, LHCII helix A, B and a CLA pair known as a potential quenching site leads to multiple residence time scales. The residence times demonstrate that the inter-pigment fluctuations occur in mostly slower time scales in the presence of multiple faster time scales comparable to conformational fluctuations of the protein and the LHCII dimer residence time. The corresponding time-dependent rates and comparable fluctuations in the LHCII manifest the emergence of dynamic disorder in LHCII. Our results lay the foundation for exploring the role of dynamic disorder in light-harvesting and heat dissipation mechanisms for future study.

## Simulation Details

CG molecular dynamic simulations of the plant thylakoid membranes are performed by using the Martini-2.2 force field. ^39,40^ The initial configuration of the thylakoid is sourced from the MARTINI website (http://cgmartini.nl/index.php/all-files-needed-for-the-simulation-of-a-thylakoid-membrane) which comprises seven distinct lipid types (see Figure S1 of supplemnetry information (SI)) and a total of 2044 lipids, 748 Na^+^, and 136 Cl^−^ molecules. The lipids in thylakoid include two types of phosphatidylglycerol: 16:1(3t)-16:0 PG, 16:1(3t)-18:3 (9,12,15) PT and two types of digalactosyldiacylglycerol: 18:3 (9,12,15)-16:0 DGDG, di18:3 (9,12,15) DGDT. In addition, the membrane contains two types of monogalactosyldiacylglycerol: 18:3 (9,12,15)-16:0 MGDG, di18:3 (9,12,15) MGDT, and sulfoquinovosyldiacyl-glycerol: 18:3 (9,12,15)-16:0 SQDG. Among the seven types of lipids, DGDG, MGDG, PG, and SQDG contain less number of the double bonds in their tails (referred to as least unsaturated lipids) compared to the PT, DGDT, and MGDT, which include a higher number of double bonds (referred as highest unsaturated lipids). Thylakoid bilayers with 64 and 128 CLA are pre-equilibrated for 1 *µ*s and 9 − 10 *µ*s using M3^22^ and Martini model^41^ respectively in previous studies. These pre-equilibrated configurations have been used as the initial configurations of the current investigations for both the M3 and Martini models. The method of inserting CLA into the plant thylakoid membrane and their simulation details are mentioned in ref^23^ in detail. The bilayers with M3 model^22^ and Martini-2.2 model^38^ of CLA are referred to as 64 M3, 128 M3 and 64 Martini, 128 Martini, respectively, in Table 1. The numbers of CLA in thylakoid are chosen in such a way that the CLA-to-lipid ratio falls below 1250, which allows enhanced Fluorescence quenching due to the CLA aggregation. ^22^ To analyze the dynamical properties of CLA aggregation, a 100 ns NVT production run is carried out with 2 fs and 10 fs time steps for M3 and Martini model, respectively, for each bilayer with 64 − 128 CLA (see Table 1).

**Table 1:**
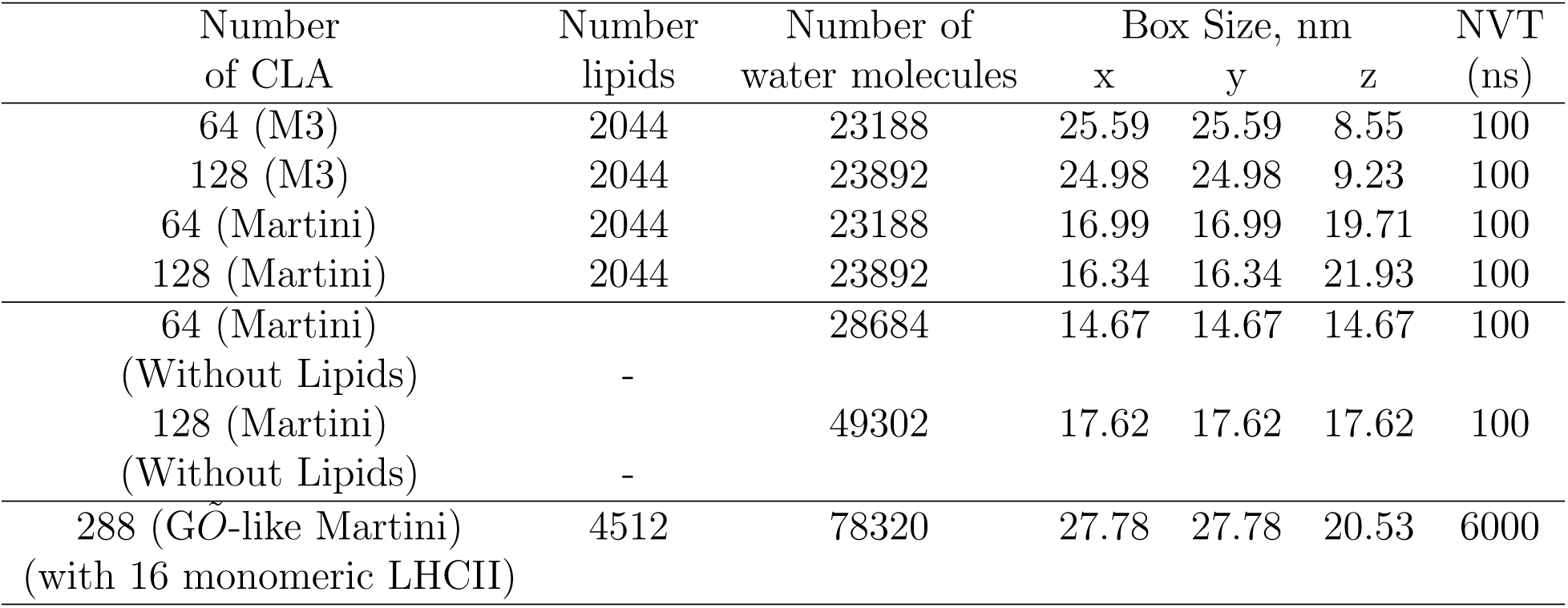
Details of systems simulated.

To show how slow fluctuations of lipids affect the residence time of CLA dimers, bilayers with 64 CLA and 128 CLA have been simulated without plant thylakoid in water solvent beads following the reference^21^ using the Martini-2.2 force field. The systems are energy minimized and a short NVT 100 ps has been performed using a v-rescale thermostat at 293 K. After that systems are equilibrated enough for 1 *µs* using an NPT ensemble with Brendsen barostat. To further analyze the dynamical properties of CLA aggregation, a 100 ns NVT production run is carried out with 10 fs time steps. The temperature is kept constant at 293 K by using a V-rescale thermostat^42^ with a coupling constant of 2 and 1 ps in the M3 and Martini models, respectively. Long-range interactions are corrected using the particle mesh Ewald method in M3 and the reaction-field method in the MARTINI model.

To study monomeric LHCII aggregations in the plant thylakoid membranes, 16 units of LHCII monomer are simulated in the presence of thylakoid bilayers with 40% non-bilayer lipids. The crystal structure of LHCII (1RWT.pdb)^1^ has the protein trimer, chlorophyll-a, chlorophyll-b, lutein, neoxanthin, and violoxanthin molecules. The AA protein monomer from the crystal structure is mapped into a CG protein configuration with the G*Õ*-like model^43^ of Martini-3.0^44^ using the program Martinize.^45^ Since dynamic disorder of the protein is associated with structural fluctuations of the protein and G*Õ*-like model can capture the conformational fluctuations of the protein,^43,46^ Martini-3.0 is used for the simulations of LHCII in the membrane. The AA co-factors of LHCII are mapped into CG representation using pycgtool^47^ following mapping schemes of Martini. ^48^ The thylakoid lipids are mapped in a similar line to Martini-3.0 as mentioned in reference.^49^ The LHCII monomer is embedded in the plant thylakoid membrane using the Python tool insane^50^ and the simulation box is replicated four times in each of the x-y directions which creates 16 units of the LHCII monomers in 4512 lipids by maintaining plant thylakoid lipid compositions. The thylakoid bilayer with 16 LHCII monomers is energy minimized followed by 100 ps short NVT. For all backbone beads of LHCII, a position restraint of 1000 kJ mol^−1^ nm^−2^ is applied for a 10 ns NPT run, and then the position restraint is reduced to half for another 10 ns NPT run. Finally, the position restraint is removed and a 5 *µ*s NPT run is carried out with semi-isotropic pressure coupling. After the NPT run, a long 6 *µ*s NVT run is carried out with 100 ps saving frequency to analyze the properties related to dynamic disorder in the aggregated LHCII monomers in the thylakoid membrane.

The LINCS algorithm^51^ is used to constrain all bonds in all systems simulated as mentioned in Table 1. In all three (XYZ) directions, periodic boundary conditions (PBC) are used. All simulations are carried out using the GROMACS 2018.1 software package.^52^

## Results and Discussion

### CLA and LHCII monomer aggregation in thylakoid

As found earlier,^22,23^ CLA dynamically forms aggregates of dimer, trimer, and tetramer when embedded in thylakoid using the M3 model. As the concentration of CLA increases, the probability of aggregation and the aggregate size increase. ^23^ Similar to the M3 model, the probability and aggregate size of CLA increase with increasing concentration of CLA using the Martini model.^41^ CLA is a smaller molecule compared to the proteins in photosynthetic machinery; the aggregation behavior of CLA occurs on a smaller time scale than the time scale of the protein’s conformational fluctuations. Since the M3 model works with a smaller time step (2 fs) compared to that used in Martini (10 fs), the M3 model is more suitable for capturing the dynamical phenomena that occur on a shorter time scale. On the other hand, Martini can achieve longer run lengths and is more suitable for capturing long-timescale behaviors. Thus, both models are employed to investigate the aggregation behavior of CLA. The lamellar thylakoid undergoes a phase transition to an inverted hexagonal phase (H*_II_*) while using the Martini model at 293 K only when the lipid:CLA ratio is below 128 : 1. ^41^ Similar L*_α_* to H*_II_* phase transformation is observed in the experimental study on CLA in dipalmitoleoyl PE (DPoPE) bilayer at 290 K when the lipid:CLA ratio is 100 : 1. ^53^ However, during the observation period of our simulations using the M3 model, thylakoid membranes remain in a lamellar phase even when the lipid:CLA ratio is below 128 : 1 and no non-lamellar phase is observed. This is attributed to the fact that using the M3 model, the time scale needed to sample the CLA-induced phase transition is not achieved. Thus, the dynamics of CLA aggregation are investigated for the lamellar thylakoid before the lamellar to non-lamellar phase transition for the M3 model. Aggregation of CLA is investigated for the non-lamellar phase of thylakoid using the Martini model. Earlier reports demonstrate thylakoid polymorphism in the absence of CLA only at high light conditions (323 K)^49^ or at lower hydration at 293 K,^54^ or above a critical concentration of CLA at 293 K^41^ suggesting critical roles of temperature, CLA concentration and hydration in facilitating phase transitions in thylakoid bilayers. As the focus of the current investigation is to understand the dynamics of CLA aggregation, not the phase behavior of lipids, CLA aggregations are compared using M3 and Martini models in lamellar and non-lamellar thylakoids, respectively.

The convergence in area per lipid confirms the equilibration of the lamellar phase for M3 and the non-lamellar phase for Martini CLA (data not shown). The CLA molecules dynamically aggregate in the bilayer, and the lipids with the lowest degree of unsaturation (DGDG, MGDG, PG, and SQDG as seen in Figure S1 of SI) are more strongly attracted to CLA aggregates (see Figure 1) showing the preferential location of the unsaturated lipids near the CLA aggregate as found in earlier studies.^10,23^

**Figure 1:**
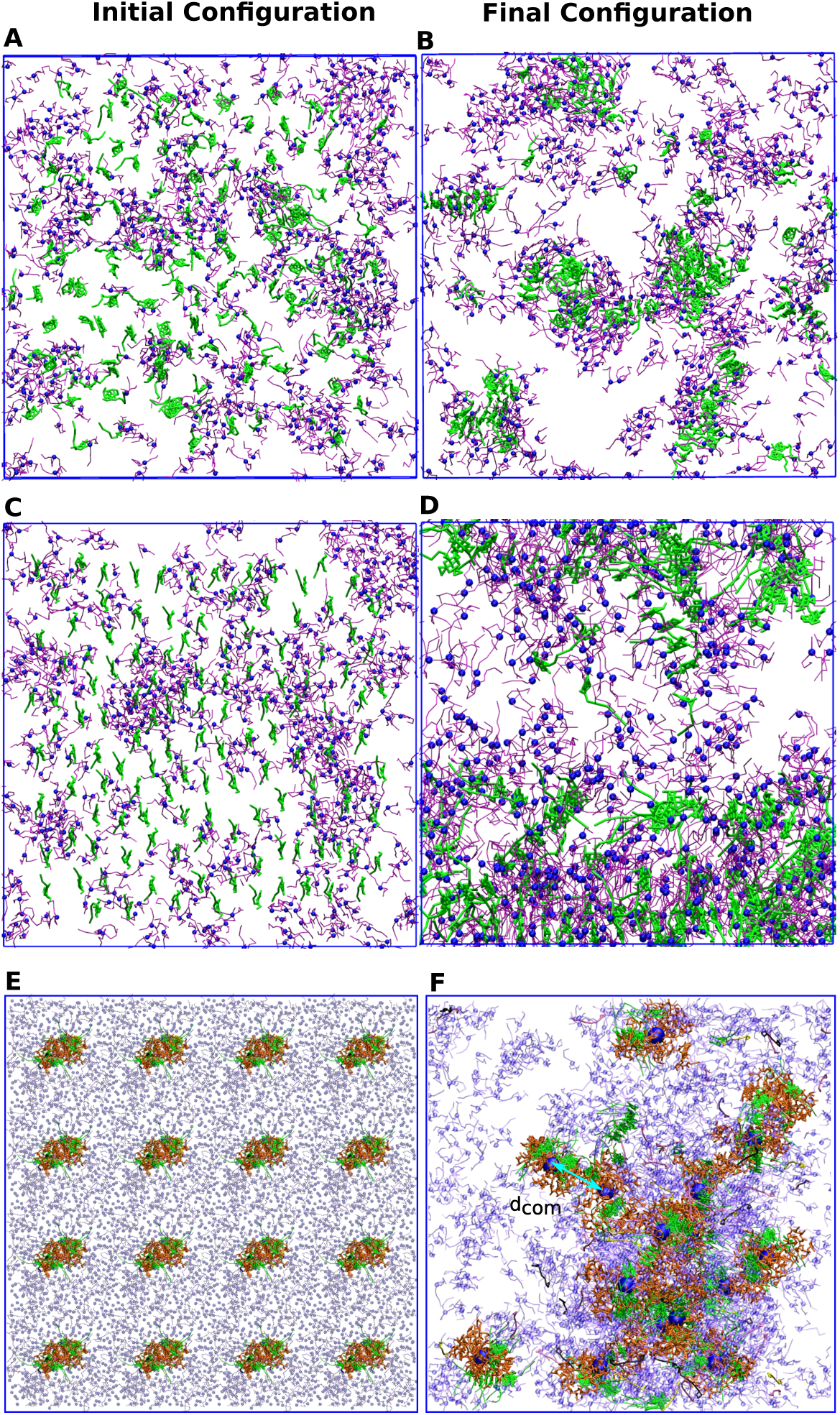
Top view of the (A, C) initial (energy minimized) and (B, D) final (100 ns NVT production run) configurations of the bilayer with 128 CLA using (A, B) M3 and (C, D) Martini model. GL1 lipid head: Blue, least unsaturated lipids: Purple, and CLA: Green. Top view of (E) initial (energy minimized) and (F) final (6 *µ*s NVT) configuration of 16 LHCII monomer in plant thylakoid bilayer. Least unsaturated lipids: Purple (transparent), proteins: orange, chlorophyll-a -b: green, lutein: mauve, violoxanthin: yellow, neoxanthin: black. The blue spheres in (F) represent the center of mass (COM) of the LHCII monomer and the arrow (cyan) represents the distance between the COM of two LHCII monomers. The remaining lipids present in the thylakoid membrane are not shown for the sake of clarity.

The initial configuration of 16 LHCII monomer embedded in the lamellar plant thylakoid membrane is shown in (Figure 1E). With time the LHCII monomers dynamically form aggregates (Figure 1E). If the center of mass (COM) of two LHCII monomers reside within the first coordination shell of 4.62 nm (location of the first peak of *g*(*r*)), they are considered to be a dimer. LHCII monomers are found to form dimer, trimer, and higher-order aggregates (see Figure S2 of SI). Upon aggregation of LHCII monomers, the thylakoid undergoes a lamellar to an inverted hexagonal phase (*H_II_*) transformation at 293 K. Similar to CLA molecules, the lipids with the lowest degree of unsaturation (DGDG, MGDG, PG, and SQDG as seen in Figure S1 of SI) preferentially locate near the LHCII aggregates (see Figure 1F).

### Contact lifetime and Waiting time of CLA dimer

To understand the dynamical behavior of CLA aggregation in plant thylakoid membranes, the contact lifetime and their waiting time for dimer formation are estimated. Our earlier study^23^ reports the average contact lifetime of each pair but does not analyze their distributions. The location of the first peak of the radial distribution function (RDF) is considered to be the cut-off distance for the dimer formation. If the distance between the COM of two CLA molecules is less than or equal to the cut-off distance, the CLA pair is considered to be in contact or as a dimer. If the distance between the COM of two CLA molecules is greater than the cutoff distance, the dimer is considered as two CLA monomers (out of contact). The dimer forming and breaking times of CLA dimer are defined as the times at which the two monomers come into contact and go out of contact.

When a CLA comes within the first coordination shell of another CLA for the first time point in the trajectory, it is considered as the first dimer formation time (*T*_1_*_f_*, Figure 2 a). The time when the CLA goes away from the cutoff distance of the partner for the first time is regarded as the first breaking time (*T*_1_*_b_*). The difference between the first breaking time and first forming times (*T*_1_*_b_* − *T*_1_*_f_*) represents the first contact lifetime (*L*_1_) of the dimer (see Figure 2 a). Similarly, the second contact lifetime (*L*_2_) is calculated from the difference between the second breaking time and second forming times (*T*_2_*_b_* − *T*_2_*_f_*) of the dimer. The aforementioned procedure is repeated throughout the trajectory for all possible pairs of dimers to compute all subsequent contact lifetimes. All contact lifetimes from all dimers across the entire trajectory have been computed, creating a comprehensive dataset of all contact lifetimes. This dataset is then used to construct a contact lifetime distribution. The contact lifetime distributions of dimers for the bilayers with 64 and 128 CLA using M3 and Martini models are shown in Figures 2 (B and C) and (D and E), respectively. The contact lifetime exhibits distributions of three to five time-scales using the M3 model. Two to three characteristic time-scales are obtained for the contact lifetime distributions of CLA using the Martini model. The probability distribution of contact lifetime is the highest for the faster time scale of 10^−1^ ps for both M3 and Martini model for 64 and 128 CLA, and probability decreases with increasing time scale. The contact lifetime distributions of 64 and 128 CLA using the Martini model exhibit two to three characteristic time scales. However, the probability of contact lifetime and waiting time distribution is less in the case of Martini than in the M3 model of CLA, suggesting that CLA dimers are more stable in the M3 model and show well-captured faster time-scale dynamics due to a smaller time step (2 fs). This result is supported by the more negative binding free energy of CLA dimer in the M3 model than in the Martini model.^10^ The emergence of multiple time scales in the contact lifetime distribution indicates the presence of dynamic disorder.

**Figure 2:**
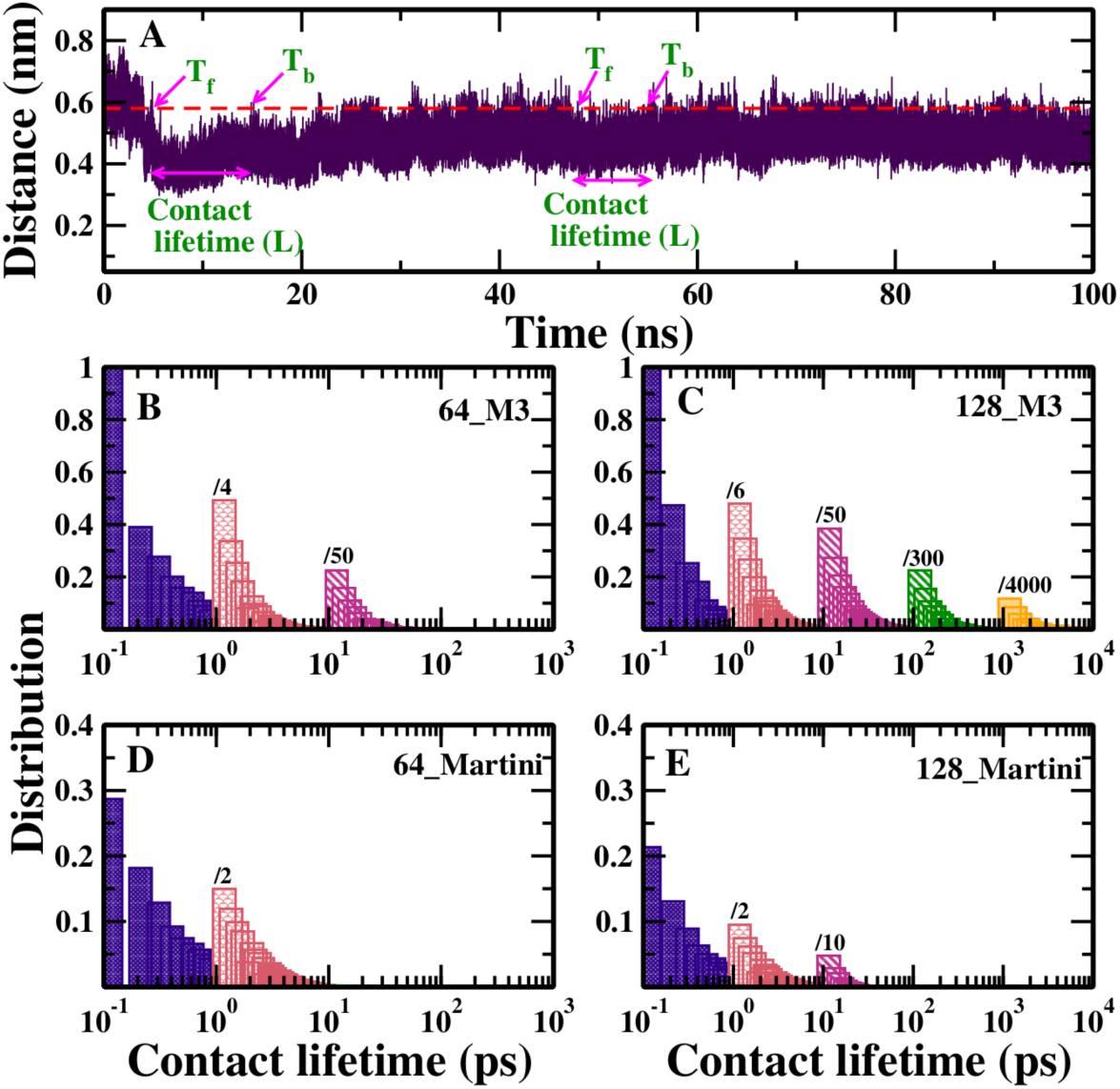
(A) Distance between two CLA COM showing a representative contact lifetime (*L*). red dotted line: cut-off distance considered for dimer formation. *T_f_* : Dimer forming time and *T_b_*: Dimer breaking time. Contact lifetime distribution of (B, C) M3 and (D, E) Martini CLA dimer for bilayer with (B, D) 64 and (C, E) 128 CLA. The number mentioned on each bar represents the multiplication factor of the probability distribution for a better comparison.

Similar to contact lifetime distributions, waiting time distributions are calculated from the waiting time for all dimers obtained during the simulation. The first waiting time of the dimer (*W*_1_) is calculated from the difference between the second forming time (*T*_2_*_f_*) and the first braking times (*T*_1_*_b_*). Figure 3 A) shows a representative waiting time (*W*). Similarly, the subsequent waiting times are calculated by identifying the breaking and next forming times. The waiting time distributions of dimers for the bilayers with 64 and 128 CLA are shown in Figure 3 (B and C) for the M3 model and in Figure 3 (D and E) for Martini model. Similar to contact lifetime distributions, the waiting time distribution of CLA dimer exhibits three to five time scales for the bilayer with 64 and 128 CLA, respectively, using the M3 model. The probability distribution of the fastest time scale (10^−1^ ps) is higher than the slowest time scale. This indicates the probability of CLA dimer going out of contact for a longer time-scale (10^2^ or 10^3^ ps) is very low. The multiple time scales in contact lifetime and waiting time distribution indicate the presence of dynamic disorder in CLA aggregation. Similar to the M3 model of CLA, the Martini CLA also shows multiple time scales (Figure 3 D and E) in waiting time distribution, suggesting the existence of dynamic disorder in the aggregation of CLA. Very similar to the M3 model, the probability of the fastest time scale distribution from waiting time is higher than the slowest time scale. Both CLA models show significantly lower but non-zero probability distributions of the slowest contact lifetime and waiting time. This indicates that the mechanism of dimer formation and breaking is mainly governed by faster dynamics with low contributions of slower dynamics. Thus, CLA aggregation occurs mainly under a fast fluctuation limit with a non-zero contribution from slow fluctuations.

**Figure 3:**
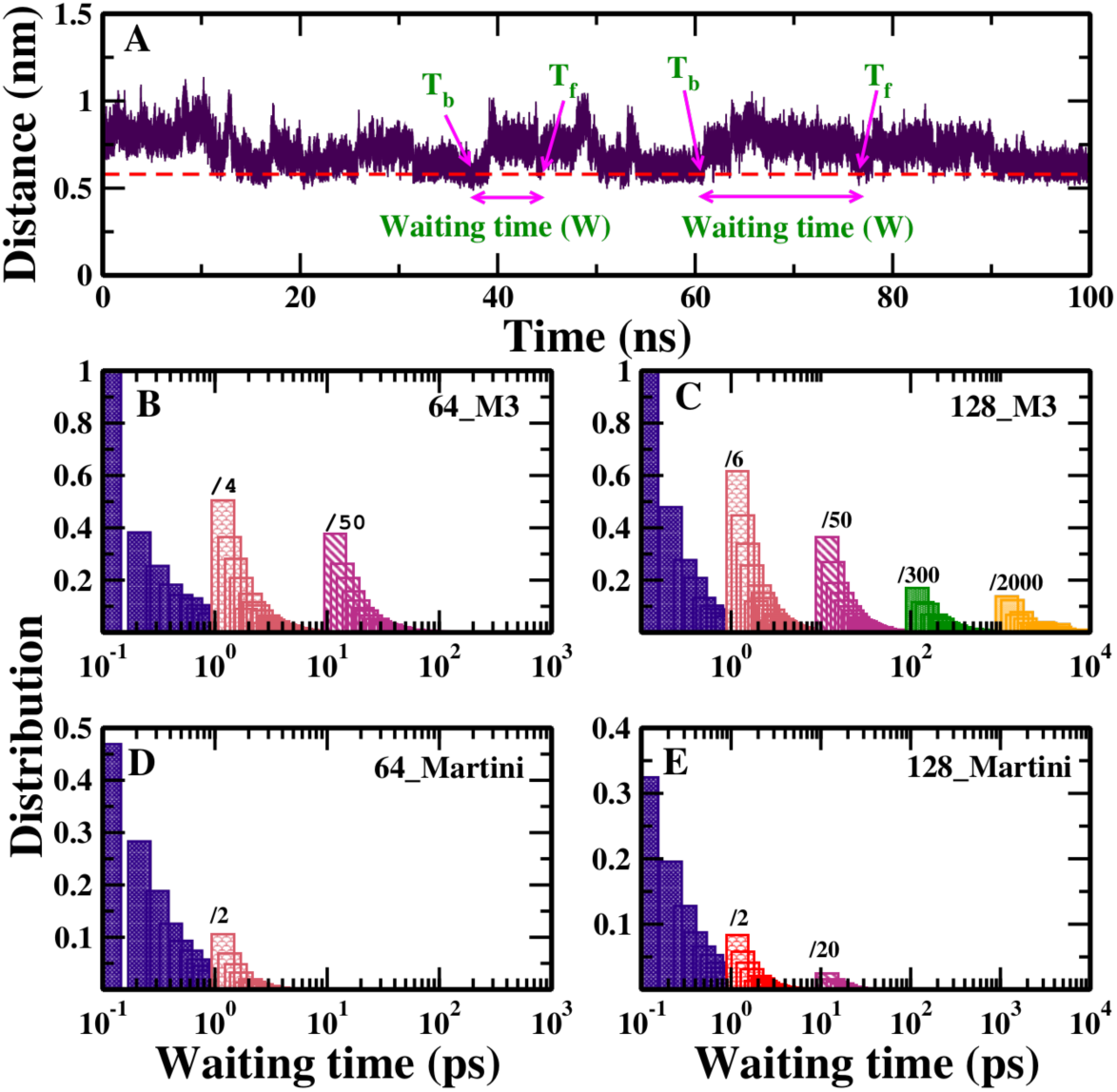
(A) Distance between two CLA COM showing a representative waiting time (*W*). Red dotted line: cut-off distance is considered for dimers out of contact. *T_f_* : Dimer forming time and *T_b_*: Dimer breaking time. Waiting time distribution of (B, C) M3 and (D, E) Martini CLA dimer for bilayer with (B, D) 64 and (C, E) 128 CLA. The number mentioned on each bar represents the multiplication factor of the probability distribution for a better comparison.

### Dynamic Disorder

A chemical reaction or a physical process involving a rate constant as a random or stochastic function of time is considered to be in a state of dynamic disorder.^55^ A conventional rate equation describing the concentration *C* of a particular species undergoing a chemical or physical process can be expressed as:

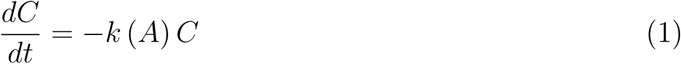

where *k* denotes the rate constant, which is a function of a control variable, *A*, which is also a function of time. When *A* is a stochastic function that varies with time, denoted as *A*(*t*), it results in dynamical disorder. The solution to the rate equation (1) that is dependent on time can be written as:

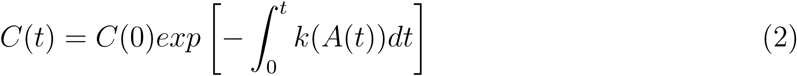

The concentration can be elucidated in terms of survival probability, *S*(*t*), of a CLA dimer that will remain or survive its state as a dimer during time *t*. Thus *S*(*t*) can be written as:^56^

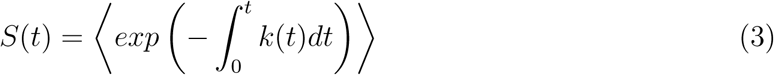

where *k*(*t*) represents the rate constant. If *k*(*t*) fluctuates very rapidly, then *S*(*t*) can be approximated by eq.4, which can be represented using a single-exponential function.

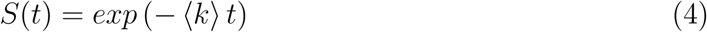

*S*(*t*) is defined by a multiexponential function when k(t) changes very slowly.

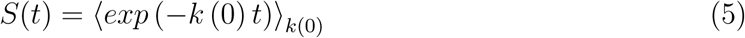

Performing the averaging in eq.3 in a direct approach is challenging. In our earlier work, to overcome this problem, Jensen’s inequality (i.e.,⟨*e*^−^*^x^*⟩ ≥ *e*^−⟨^*^x^*^⟩^) is used for exponential function where both upper and lower bounds for the survival probability have been derived in time domain.^57^ This is achieved by employing a trial action that allows for accurate solvability of the path integrals.

Dynamic disorder refers to time-dependent fluctuations in a reaction rate when a molecule experiences different conformations in the free energy landscape due to temporal variations in the control variable, *A*. If these fluctuations are fast as mentioned in equation (4), the reaction rate changes rapidly, they average out over time, leading to a single effective rate constant and resulting in exponential kinetics.^34,58^ Static disorder describes slow fluctuations of the control variable, where changes remain effectively constant over the relevant timescale of observation, and is also described by a non-exponential function.^34,58^

### Survival probability and Rate of CLA

The survival probability of CLA dimer is determined using the following equation:^59^

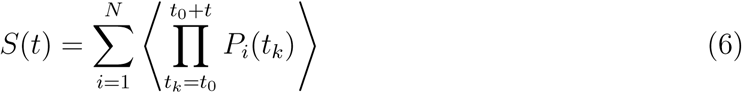

Where *S*(*t*) is the probability of finding a CLA dimer within the cutoff distance (first rdf peak) at time *t*, *N* denotes the total number of molecules, and the angular brackets denote averaging over the time origins *t*_0_. *P_i_*(*t*) = 1 if the CLA dimer is present within the cutoff at time *t*, and *P_i_*(*t*) = 0 otherwise. *S*(*t*) is calculated after averaging over five uncorrelated datasets and shown in Figure 4A-B. The decay of the *S*(*t*) of 128 CLA molecules is slower than that of the 64 CLA in both M3 and Martini models (Figure 4A and B). The dimer survival probability of 128 CLA in the M3 model does not decay completely within 1 ns (see figure 4 (A)), unlike other cases indicating the survival of a dimer for a longer time scale. Residence time scales for the CLA dimer have been extracted from a tri or five-exponential function fitting of the survival probability average over five data sets using the following equation:

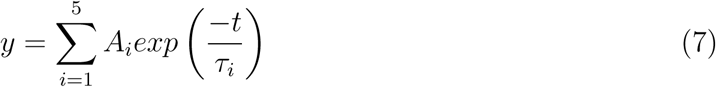

**Figure 4:**
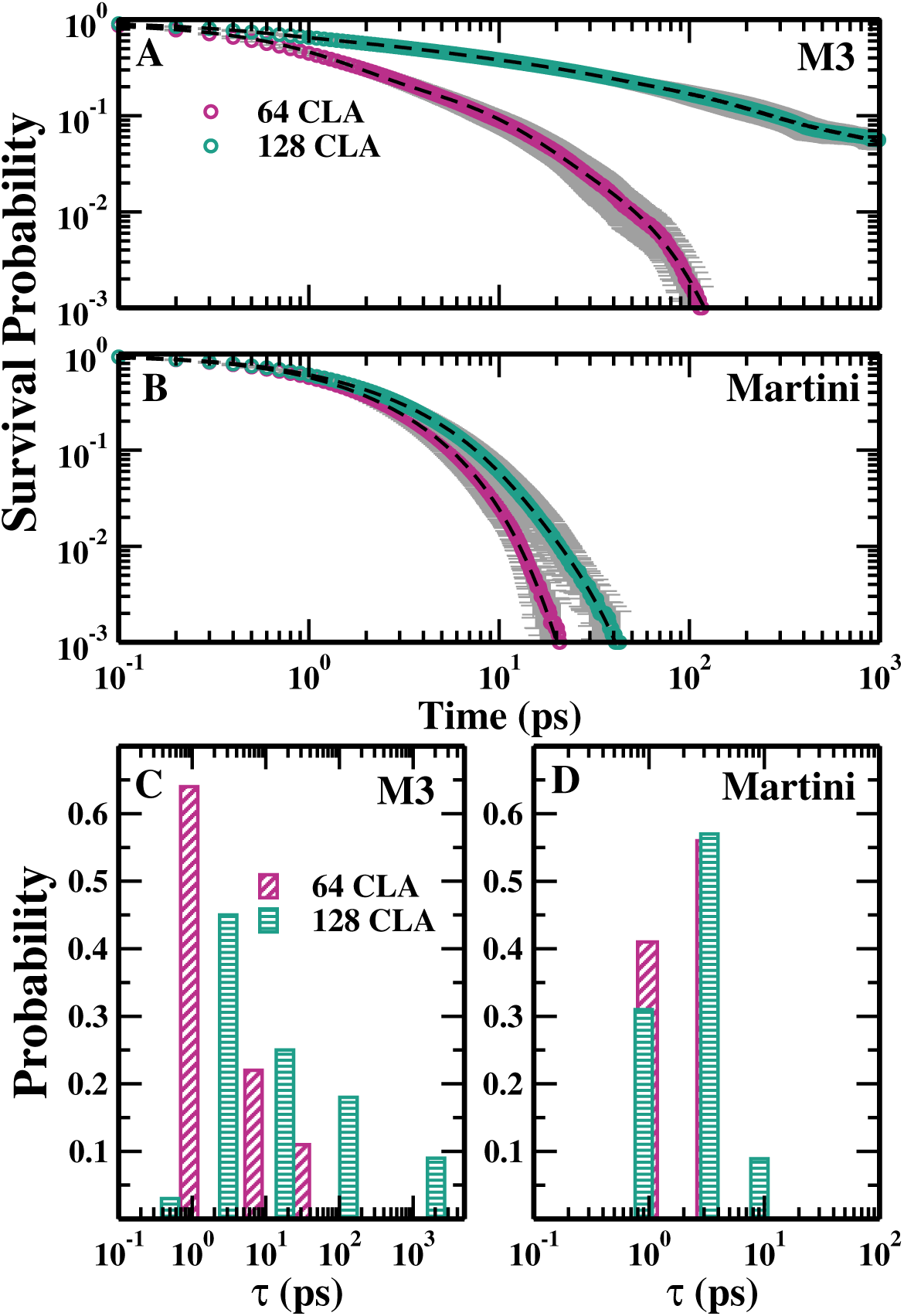
Survival probability of CLA dimer formation of (A) M3 (B) Martini CLA for the bilayer with 64 and 128 CLA.Multiple residence time scales of the CLA dimer for (C) M3 and (D) Martini CLA for the thylakoid with 64 and 128 CLA.

The timescales obtained from the fitting are shown in Figure 4C-D. CLA dimer exhibits three and five characteristic time scales for 64 and 128 CLA, respectively, using the M3 model (see Figure 4 (C)) and two and three characteristic time scales for 64 and 128 CLA, respectively using Martini model (see Figure 4 (D)). At short times, the residence time for the 64 CLA shows a higher amplitude than that for the 128 CLA. This observation suggests a higher survival probability of the 64 CLA as compared to that of the 128 CLA at short times, while, at long times the trend is reversed. For 64 CLA, dimers form and break rapidly within a very short timeframe. The increased crowding in 128 CLA likely stabilizes CLA dimers, causing them to stay in contact for longer durations compared to dimers in the bilayer with 64 CLA. Within our simulation period, we observe that CLA molecules form dimers, break and reform the dimer in the presence of trimer, tetramer, and higher order aggregates. Therefore, the transitions between multistates occur within the observation period suggesting time-dependent dimer formation rather than states with very slow relaxation times. Both the Martini and M3 model of CLA shows that the mechanism of dimer formation is primarily governed by faster dynamics with comparatively lower but non-zero probability distributions for the slowest contact lifetimes and waiting times, suggesting that CLA aggregation occurs predominantly under a fast fluctuation limit (see Figure 4). This suggests that while slower dynamics are present they are not slower than the observation time scales to behave like fixed states, unlike the case of static disorder. This is a hallmark of dynamic disorder highlighting the presence of dynamic disorder scenarios with major fast and minor slow fluctuations for the CLA aggregation process. The multiple timescales from the survival probability agree well with those obtained from the contact lifetime. The multiple time scales obtained from survival probability, contact lifetime, and waiting time using both models strongly support the presence of dynamic disorder in CLA aggregation. Moreover, the survival probability changes as temperature changes as shown in Figure S6 of the SI. Therefore, CLA aggregation is considered to follow dynamic disorder.

To understand how slow fluctuations of lipids affect the residence time of CLA dimers, 64 CLA and 128 CLA have been simulated without plant thylakoid in water solvent beads. The probability distributions of CLA aggregates without lipids and with lipids within the first coordination shell of 0.45 nm for 64 CLA and 0.5 nm for 128 CLA show the formation of dimer, trimer, and higher order aggregates (see Figure S3 in the supplementary material (SI). Slow fluctuations and redistributions of lipids allow time for CLA molecules to find optimal orientations relative to each other. This impacts how long molecules remain in aggregated states. The comparison of the aggregation behavior of CLA in the presence and absence of lipids in thylakoid membranes reveals significant differences in their survival probability decay patterns. Figure S4 of the SI illustrates the survival probability of the CLA dimer, comparing its behavior in the presence and absence of thylakoid lipids. In the absence of lipids where CLA aggregate in water, the molecules lack the structural support of the thylakoid membrane, thus, the survival probability decays more rapidly due to increased exposure to environmental fluctuations, and fewer stabilizing pigment-lipid interactions. In contrast, when CLA aggregation occurs within the thylakoid membrane, the survival probability exhibits a slower decay. Figure S5 of the SI represents the residence time of the CLA dimer, obtained from the survival probability shown in Figure S4 of SI. The residence time of the CLA dimer is shorter in the absence of a thylakoid membrane indicating a faster dynamic in this environment. A higher probability of shorter residence time in the absence of thylakoid lipids aligns with the observed faster dynamics. This enhanced stability is attributed to the crucial structural support provided by the thylakoid membrane, stabilizing pigment-lipid interactions, and the membrane’s buffering effect against fluctuations. Thus the slow fluctuations of lipids increase the residence time of the CLA dimer.

Next, to understand the kinetics of the CLA dimer, the rate of decrease in survival probability of the CLA dimer is calculated using the following equation: ^57^

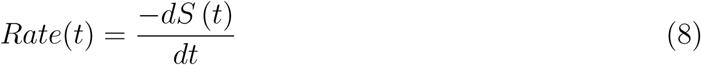

where *S*(*t*) is the survival probability. The rate is calculated from the survival probability obtained from uncorrelated five datasets and averaged, and in the presence of the thermal noise characterized by the fluctuating force *ζ* that is exerted on the surrounding beads which are primarily CG lipid beads (GL1) of thylakoid membranes in the MD simulation. Thus the noise averaging can be explicitly calculated from the MD trajectories of respective beads of interest present in the surroundings. The autocorrelation function of *ζ* ^60,61^ of GL1 lipid beads situated within 0.98 nm of the central CLA bead initially shows local valleys due to the non-uniformity in the surrounding molecules which slowly converges by 5 ps (Figure S7 A of the SI). The non-zero slope of the power spectrum (Figure S7 B of the SI) obtained from the Fourier transform of the autocorrelation function indicates colored noise on the GL1 beads on the molecular scale. Similar colored thermal noise has been found earlier for both polar and non-polar solute molecules in water. ^60,61^ As the CLA concentration increases, the dimer survival rate decays for all cases, as seen in Figure 5 (A and B). The rate follows a bi-exponential function for the M3 model and a tri-exponential function for Martini CLA. The multi-exponential behavior of the rate further supports the presence of dynamic disorder in CLA aggregation for both models of CLA.

**Figure 5:**
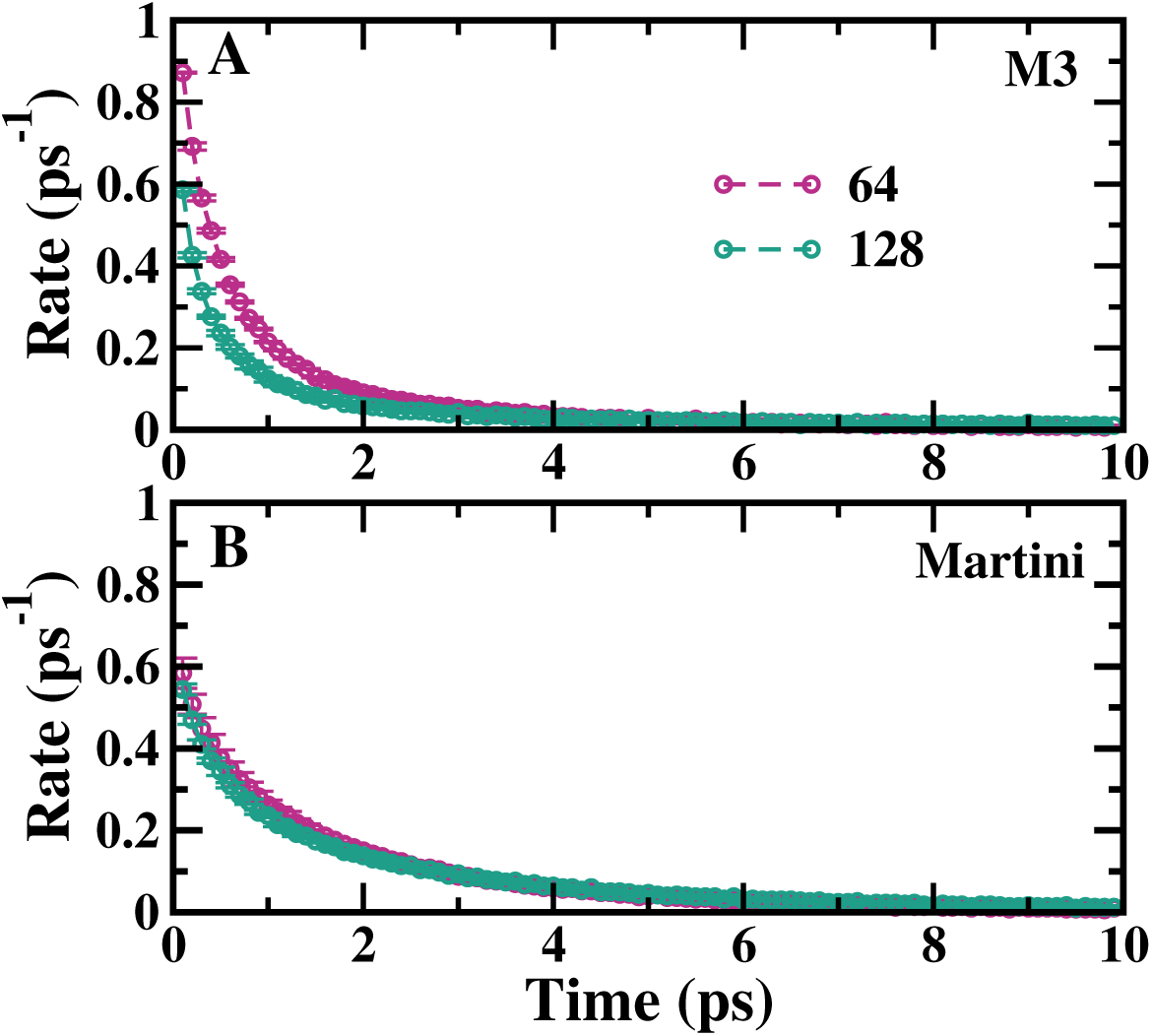
Rate of CLA dimer survival for (A) M3 and (B) Martini model of CLA for the thylakoid with 64 and 128 CLA.

### Dynamic Disorder in aggregated LHCII in thyalakoid

The aggregation of the LHCII monomers and trimers under excess illumination ^62^ can be relevant for the activation of xanthophyll cycle.^63,64^ The aggregation of the LHCII has been considered as a switch between energy dissipative and light-harvesting states.^63^ Previous MD simulations show that the N-terminal fluctuations and orientations of helix A and B of the LHCII in the membrane can cause a large variation in CLA 611-CLA 612. ^37^ Therefore, distances between these pigments and their orientations with respect to the LHCII can be the most sensitive and crucial indicators to identify any correlation between conformational fluctuations of the LHCII and inter-pigment coupling relevant to the light-harvesting or energy-dissipating states. Thus N-terminal flexibility has been estimated from the end-to-end distance (*d*) of the N-terminal of the LHCII (Figure 6 A) constructed between one of the trimerization motifs, 12 ARG (R), and its interacting partner, 34 ASP.^3^ The effect of protein conformational fluctuations on inter-pigment distances has been monitored from the end-to-end distance of the N-terminus (*d*), the distance between 611 − 612 CLA (*D*), the angle (*α*) between two helices A and B, the angle between N-terminal end-to-end vector and the vector joining 611 − 612 CLA of the corresponding monomer (Figure 6 B-C). The time evolutions of all these quantities along with the distance between two LHCII monomer COM (Figure 1F) for one pair indicate concerted fluctuations of the N-terminal to Helix A/B and 611 − 612 CLA in Figure 6 D-I. Broad distributions of all such distances and angles from all the trajectories for all the pairs (Figure 6 J-N) suggest the existence of multiple conformational substates of the aggregated LHCII and CLA.

**Figure 6:**
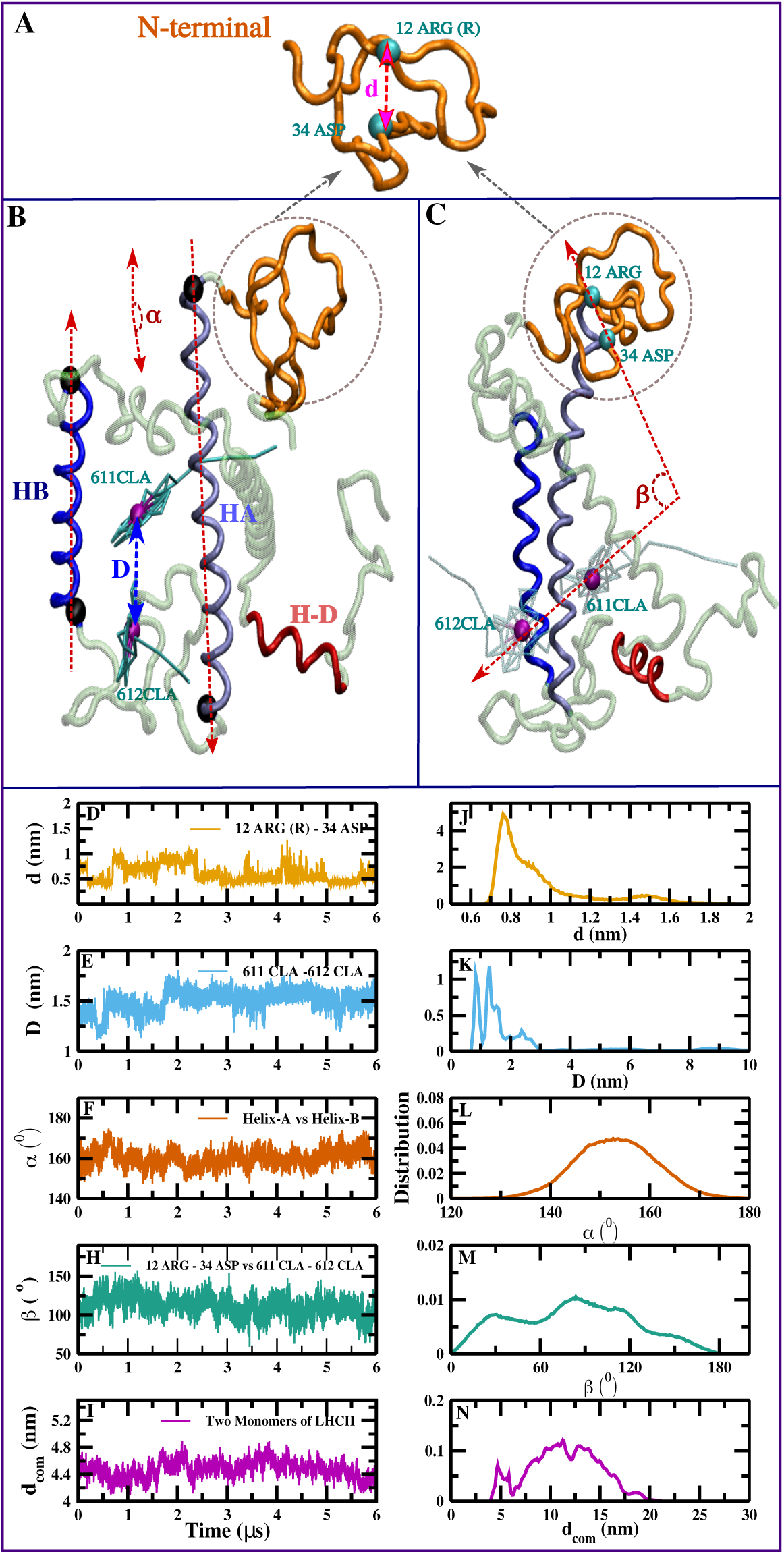
A) Snapshot of the N-terminus of the LHCII monomer showing the distance (*d*) between two interacting residues: 12 ARG (R-trimerization motif) and 34 ASP, B) LHCII monomer representing distance (*D*) between 611 − 612 CLAs and angle (*α*) between helix A and B, C) angle (*β*) between two non-bonded vectors constituted from N-terminal residues, 12 ARG-34 ASP and 611 − 612 CLAs. D)-I) Time evolution and J)-N) distribution of *d*, *D*, *α*, *β*, *d_COM_*.

The location of the most probable distribution in each case discussed in Figure 6 D-H is considered as the cutoff for calculating the respective survival probability, *S*(*t*), of *D*, *α*, and *β* as shown in Figure 7A-C. To calculate the survival probability of the LHCII dimer shown in Figure 7 D, the cutoff is chosen as the first peak of *g*(*r*) (4.62 nm) between the COM of the LHCII monomer. The decay of the *S*(*t*) of 611 − 612 CLA is the slowest suggesting that the inter-CLA distance remains conserved for long. The slower decay of the inter-CLA *S*(*t*) also slows down the *S*(*t*) of *β* which estimates the relative orientation of the N-terminal to the 611 − 612 CLA pairs. In contrast, the faster decay of *α* measuring the orientation between helix A and B suggests faster fluctuations of protein conformers. The decay of *S*(*t*) of the LCHII dimers (*d_COM_*) is similar to that of the protein conformers, *α*. *S*(*t*) in Figure 7 A-D are fitted with multi-exponential functions to extract the residence time-scales shown in Figure 7 E-H. The LCHII dimer residence time scales are comparable to the conformational fluctuations of the protein which are relatively faster than the residence time of the CLA pair. Therefore, the LHCII dimer formation and the protein conformational fluctuations can be considered as fast fluctuations. The residence time scales of the CLA pair have a slower residence time scale with high probability in addition to the time scales comparable to protein dimer formation and conformational fluctuations. The rate calculated from the *S*(*t*) for all cases is time-dependent (Figure 8). Within our simulation period, the LHCII monomers interconvert within multistates of dimer, trimer, tetramer, and higher-order aggregates. Simultaneously conformations of the protein, specifically, N-terminus, helix A and B change, which in turn affects the survival of the CLA pair responsible for the poised state of the LHCII. The same CLA pair is proposed to be the potential candidates for the quenching site. ^37^ The non-exponential decay of the *S*(*t*) and the time-dependent rate highlight the presence of dynamic disorder with fast fluctuations of protein conformations and aggregation correlated with major slow fluctuations of the pigments.

**Figure 7:**
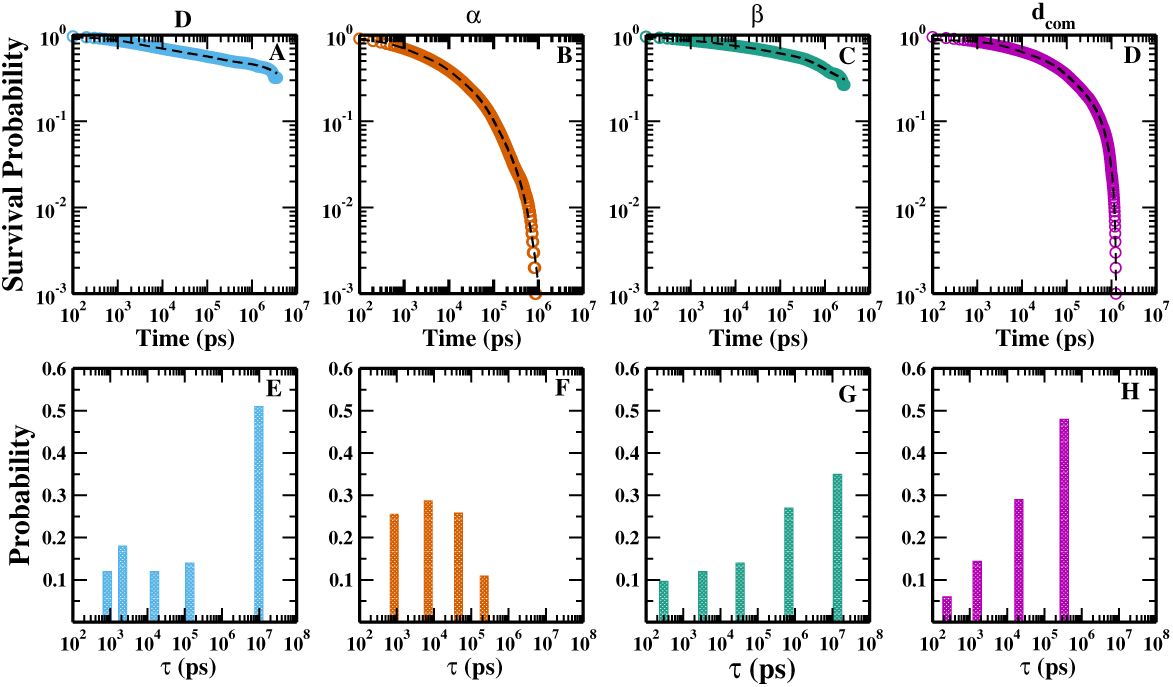
Survival probability of (A) *D*, (B) *α*, (C) *β*, and (d) *d_com_* and multiple residence time scales of (E) *D*,(F) *α* (G) *β*, and (H) *d_com_*.

**Figure 8:**
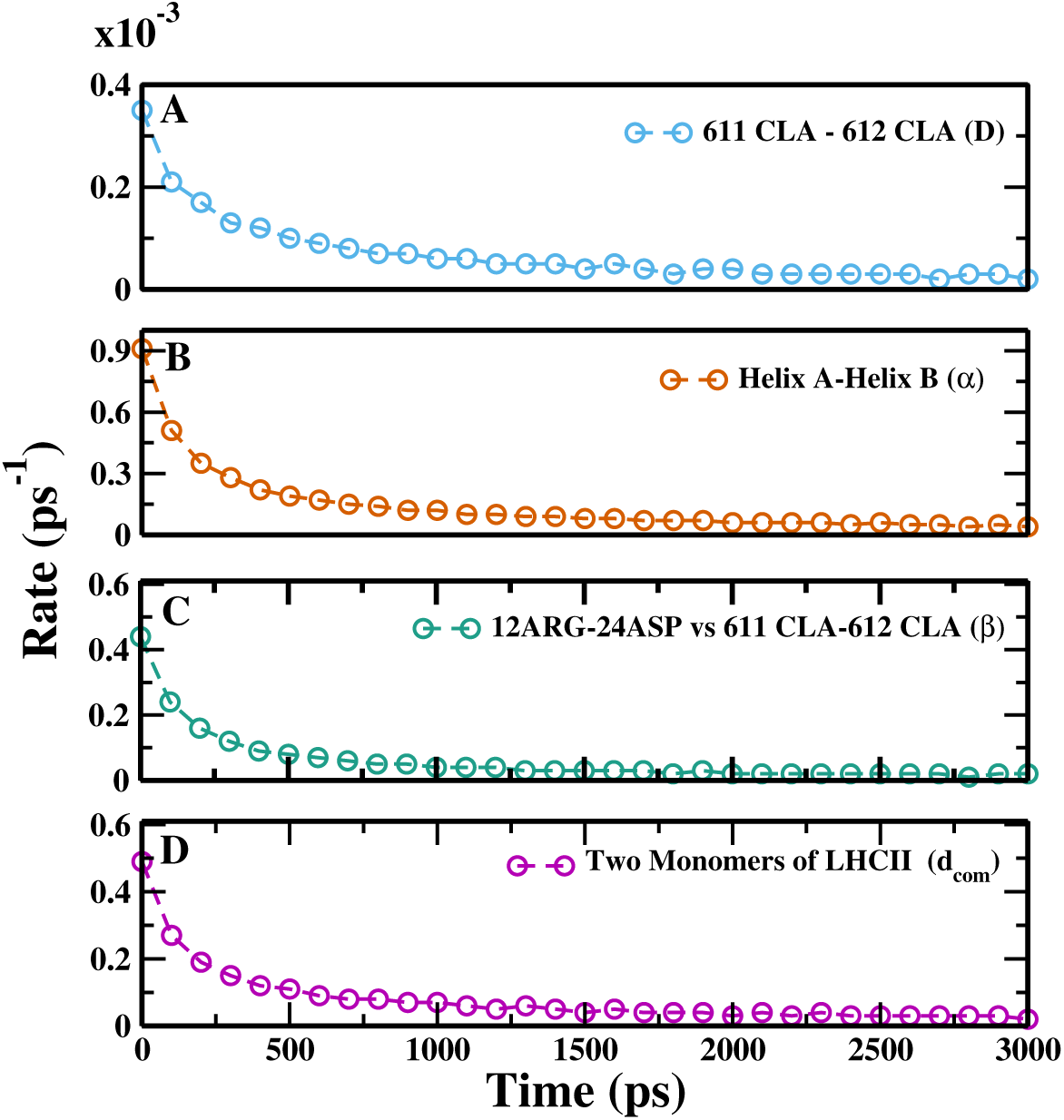
Rate of survival of (A) *D*, (B) *α*, (C) *β*, and (d) *d_com_*.

## Conclusions

To summarize, chlorophyll and LHCII aggregations in thylakoid membranes have been investigated using CG simulations where the CLA molecule is modeled by a previously derived M3 model^22^ and Martini-2.2 force-fields.^38,39^ Martini-3.0 is used for the simulations of LHCII in the membrane where the lipids are mapped following reference ^49^ and the LHCII monomer from the crystal structure (1RWT.pdb^1^) is modeled with the G*Õ*-like model^43^ of Martini-3.0^44^ to capture protein conformational fluctuations. CLA dynamically forms aggregates that break and reform throughout the trajectory in both models. The distance fluctuations of CLA molecules to form dimers occur in time scales which are dependent on the formation and breakage of higher-order aggregates in the presence of thylakoid membranes. The transitions of the dimers to trimer, tetramer, or higher-order aggregates lead to multi-state transitions which are coupled to one another and can interconvert amongst themselves at rates ranging multiple timescales. Thus CLA aggregation occurs over a broad range of time scales and exhibits intricate fast and slow fluctuations. The contact lifetime and waiting time distribution of CLA dimers show the existence of multiple time scales in both the M3 and Martini models of CLA. The probability distributions of contact lifetime and waiting time decrease with increasing time. Similar to the multiple ranges of time scales of contact lifetime and waiting time distributions, the survival probability and rate of CLA dimer follow non-exponential decay with multi-exponential relaxation times or residence times. Although the quantitative comparison of time scales can not be done between CG simulations and experiments, fluorescence quenching experiments demonstrating the existence of multiple lifetimes of CLA^65^ and aggregation^22^ validate the aggregation and multiple residence time phenomena observed in our simulations. The CLA aggregation dynamics are governed by the most populated fast fluctuations combined with less populated slow fluctuations. Such non-exponential survival probability is a manifestation of dynamic disorder. This affects the dimer residence time and makes the rate time-dependent, unlike conventional rate theories where the rate constant is independent of time. The presence of lipids in the thylakoid membrane reduces the transition rate from one state to the other due to the slow fluctuations of lipids which makes the dimer residence time slower. The dynamic disorder in CLA aggregation is especially important in the context of LHCII aggregation which is enhanced during NPQ and can be instrumental in finding energy transfer pathways under light-harvesting and NPQ conditions.^7^ The LHCII monomers embedded in thylakoid membrane dynamically form dimers and higher-order aggregates which are interconvertible among each other. Similar LHCII aggregates in membranes have been found earlier in MD simulations and NMR and fluorescence experiments.^66^ The multistate transitions are reflected in the multi-exponential decay of the LHCII dimer survival probability and time-dependent rate. The residence time scales of the LHCII dimer are similar to the conformational fluctuation time scales of the protein, specifically, the N-terminus, helix A, and B known to largely affect the inter-pigment coupling between 611 − 612 CLA, a potential quenching site.^37^ The temporal behavior of the survival probability of the 611 − 612 CLA pair is similar to that of its relative orientation to the N-terminus of the LHCII which follows a non-exponential decay and time-dependent rate. Thus the conformational fluctuations and aggregation of the LHCII occur under a fast fluctuation limit in the presence of low probable but comparable and high probable slower inter-pigment distance fluctuations highlighting the presence of dynamic disorder in LHCII. The emergence of dynamic disorder in CLA and LHCII aggregation indicates the underlying dynamics are not governed by the general transition state theory. This essentially demonstrates that the dynamics of the LHCII aggregation involve dynamic couplings of fast and slow fluctuations leading to the emergence of dynamic disorder. Thus, their reaction dynamics will be beyond the scope of a conventional free-energy picture. Since the aggregation of the LHCII is known to act as a switch between energy dissipative and light-harvesting states,^63^ comprehending the complex dynamics and disorder in CLA and LHCII aggregation can offer a profound understanding of the control mechanisms of photosynthesis and the plant’s capacity to adjust to variations in external light conditions.

## Supporting Information Available

The supplementary material provides the figures describing lipid compositions of thylakoid membranes, distributions of different orders of aggregates of LHCII in thylakoid membranes and CLA aggregates in water without membranes, the dimer survival probability and residence time without membranes, survival probability of CLA dimer at different temperatures, autocorrelation function and power spectrum of surrounding noise of CLA.

## Acknowledgement

R.S. thanks the Council of Scientific and Industrial Research (CSIR), India, for the fellowship.

## Notes

There are no conflicts to declare.

