## Supplementry Information for "Dynamic Disorder in Chlorophyll Aggregation and Light-Harvesting Complex II in the Plant Thylakoid Membranes using Coarse-Grained Simulations"

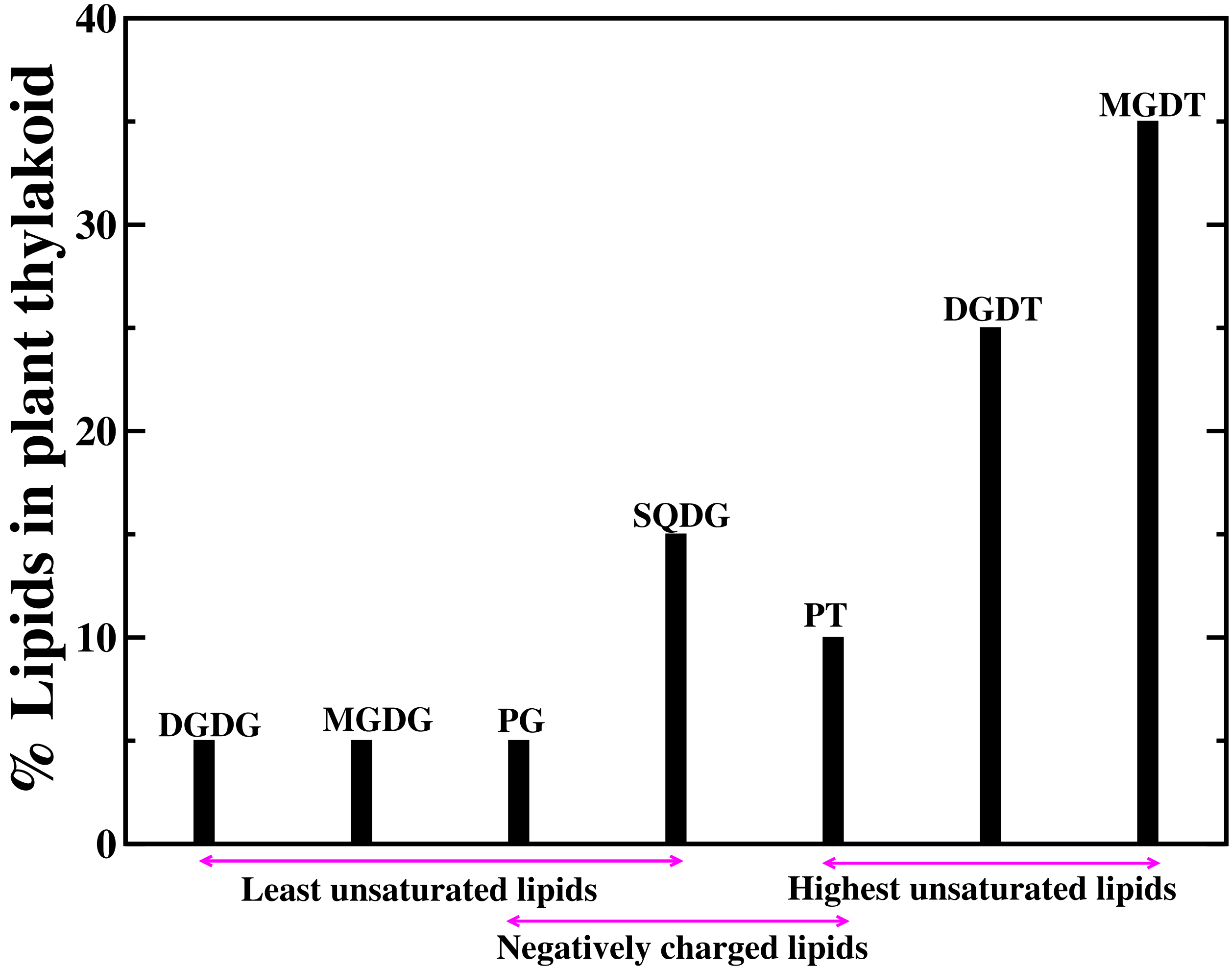


Figure S1: Lipid compositions in plant thylakoid membrane.

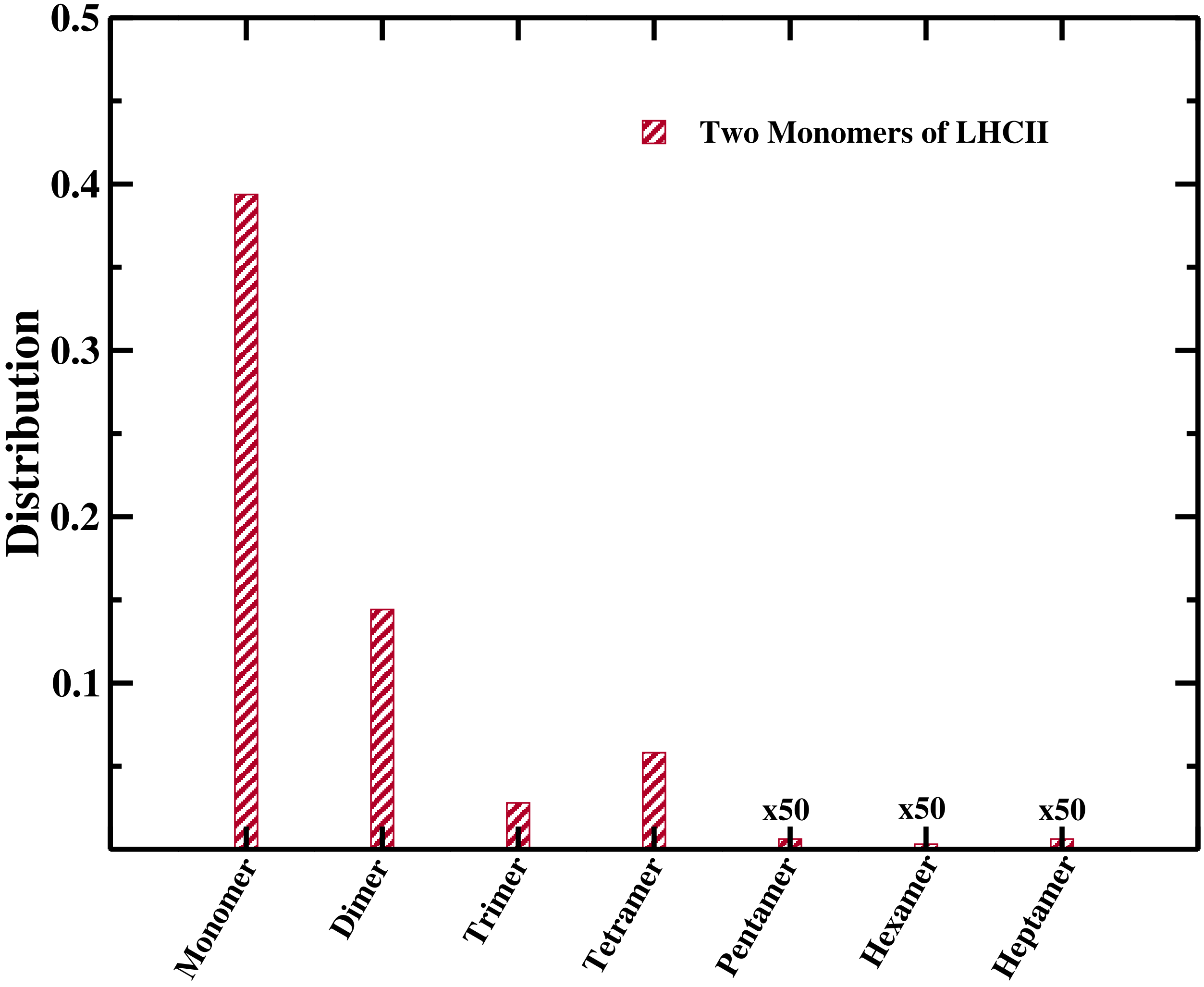


Figure S2: Probability distributions of monomer LHCII aggregates in thylakoid lipids for the first coordination shell. For clarity, the probabilities are rescaled and multiplied by different factors as shown in the plot.

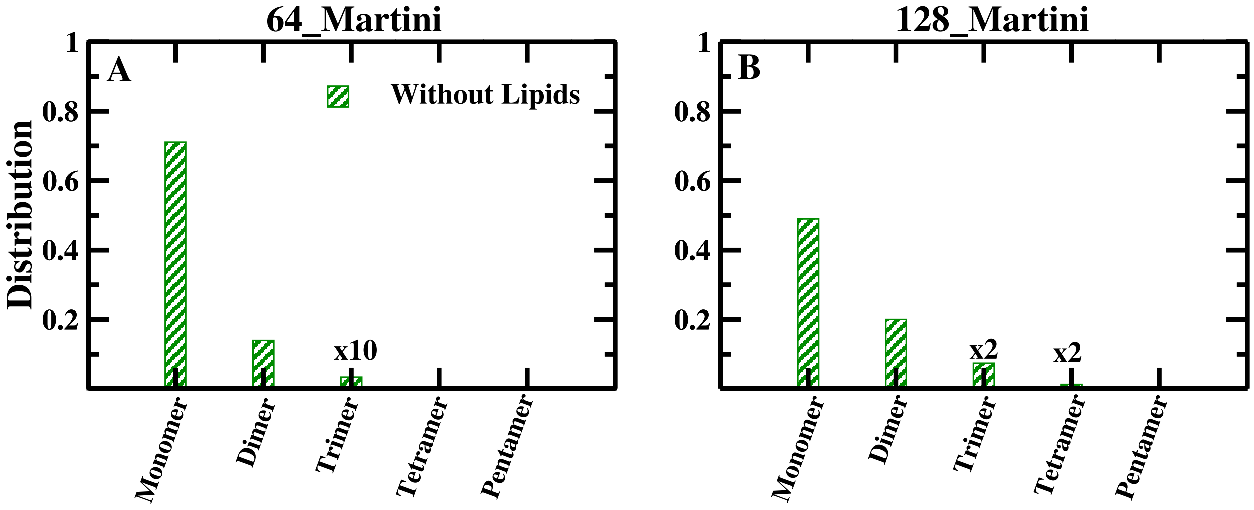


Figure S3: Probability distributions of CLA aggregates in water without lipids for the first coordination shell for A) 64, B) 128 CLA. For clarity, the probabilities are rescaled and multiplied by different factors as shown in the plot.

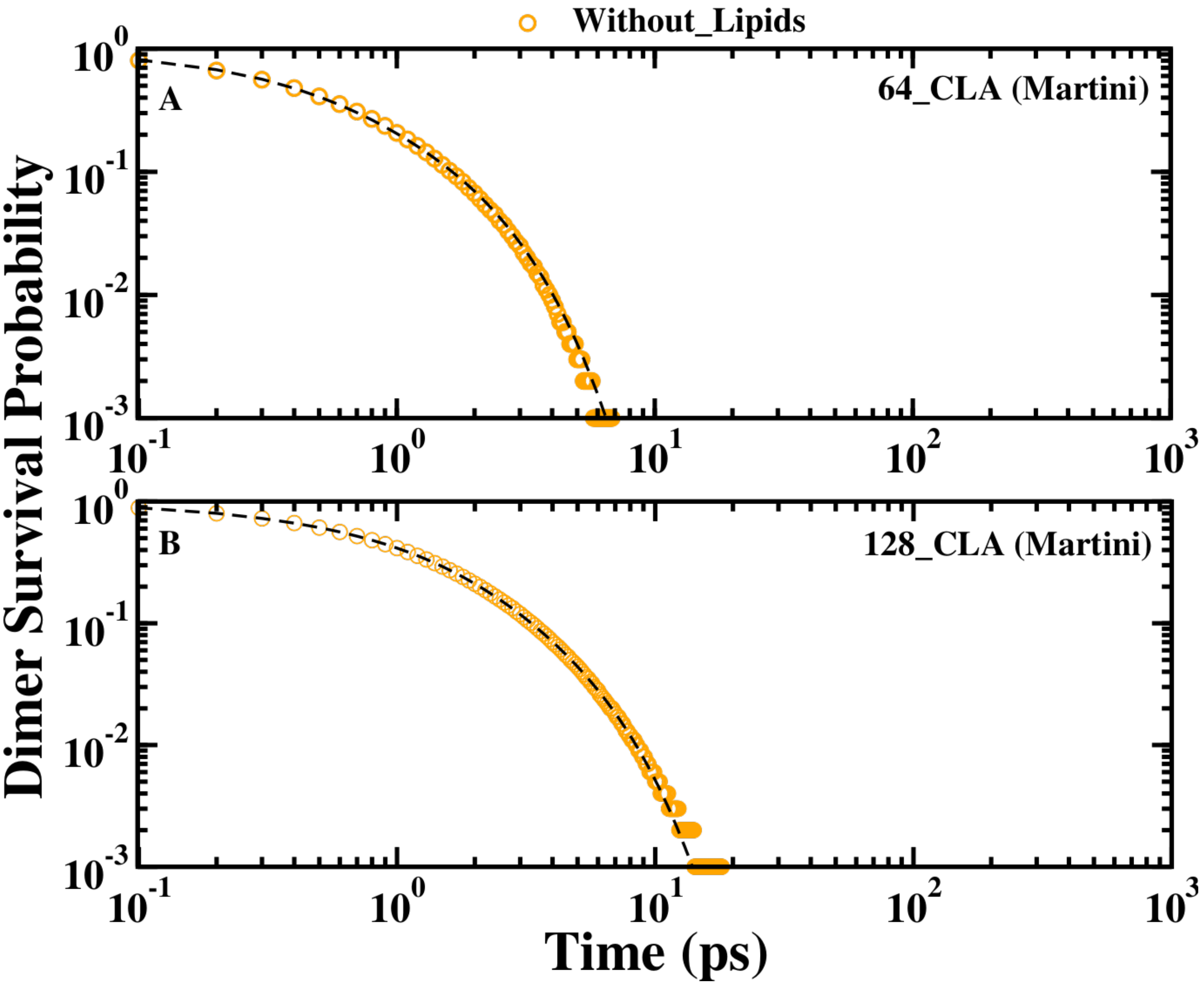


Figure S4: Survival probability of CLA dimer formation of (A) 64 CLA (B) 128 CLA without thylakoid using Martini force-field.

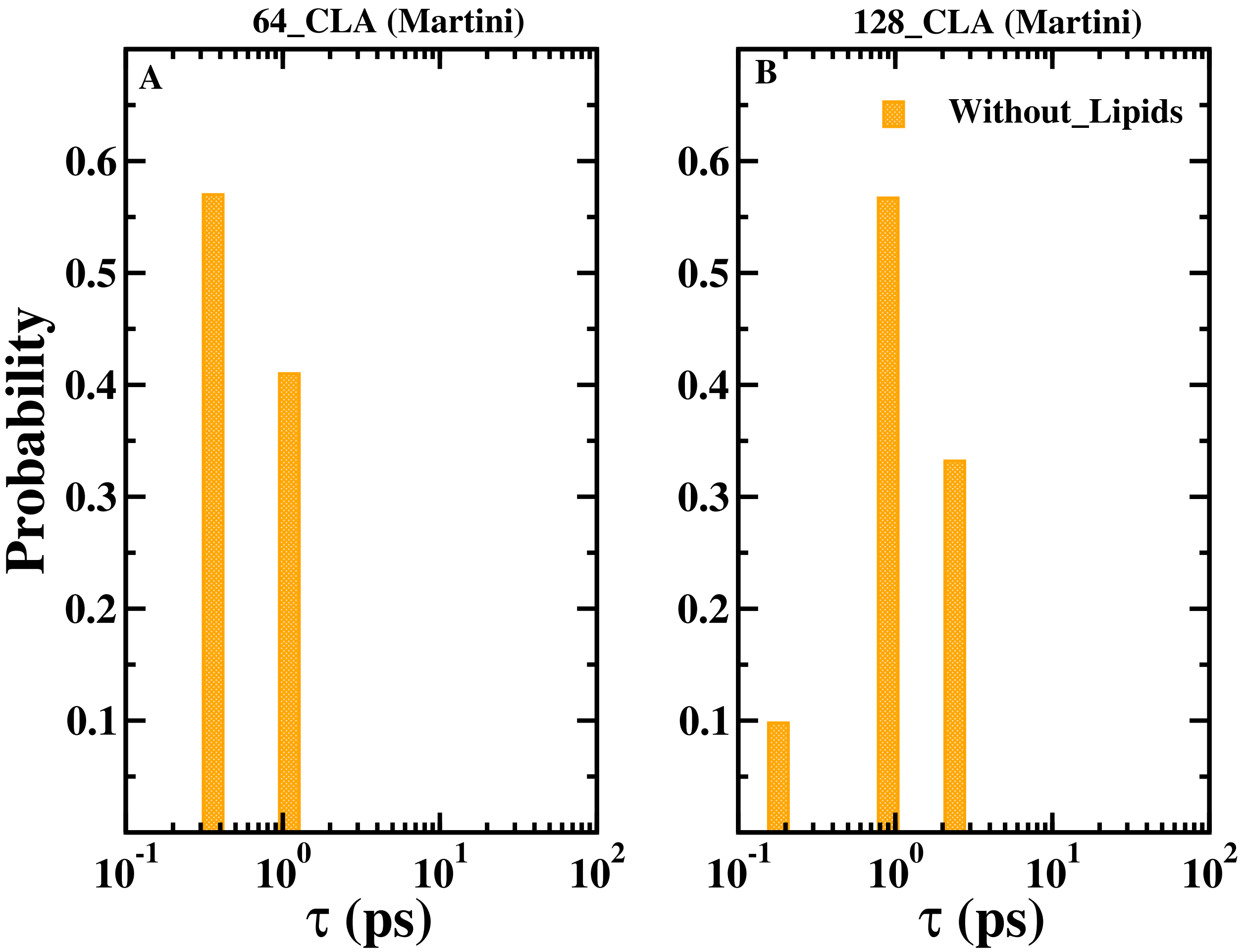


Figure S5: Multiple residence time scales of the CLA dimer for (A) 64 CLA and (B) 128 CLA without thylakoid using Martini force-field.

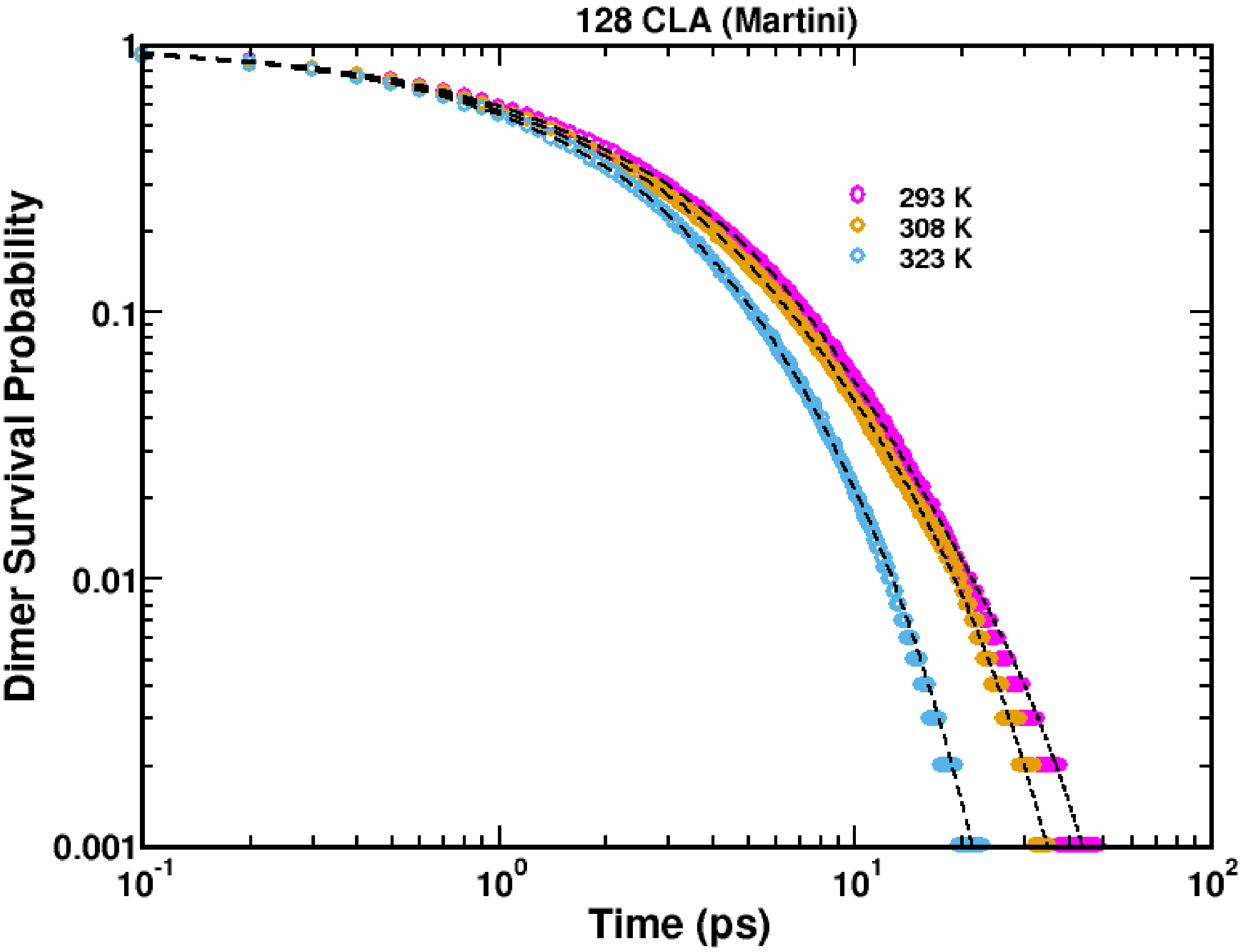


Figure S6: Survival probability of CLA dimer formation of Martini for 128 CLA in thylakoid at three temperatures, 293 K, 308 K and 323 K.

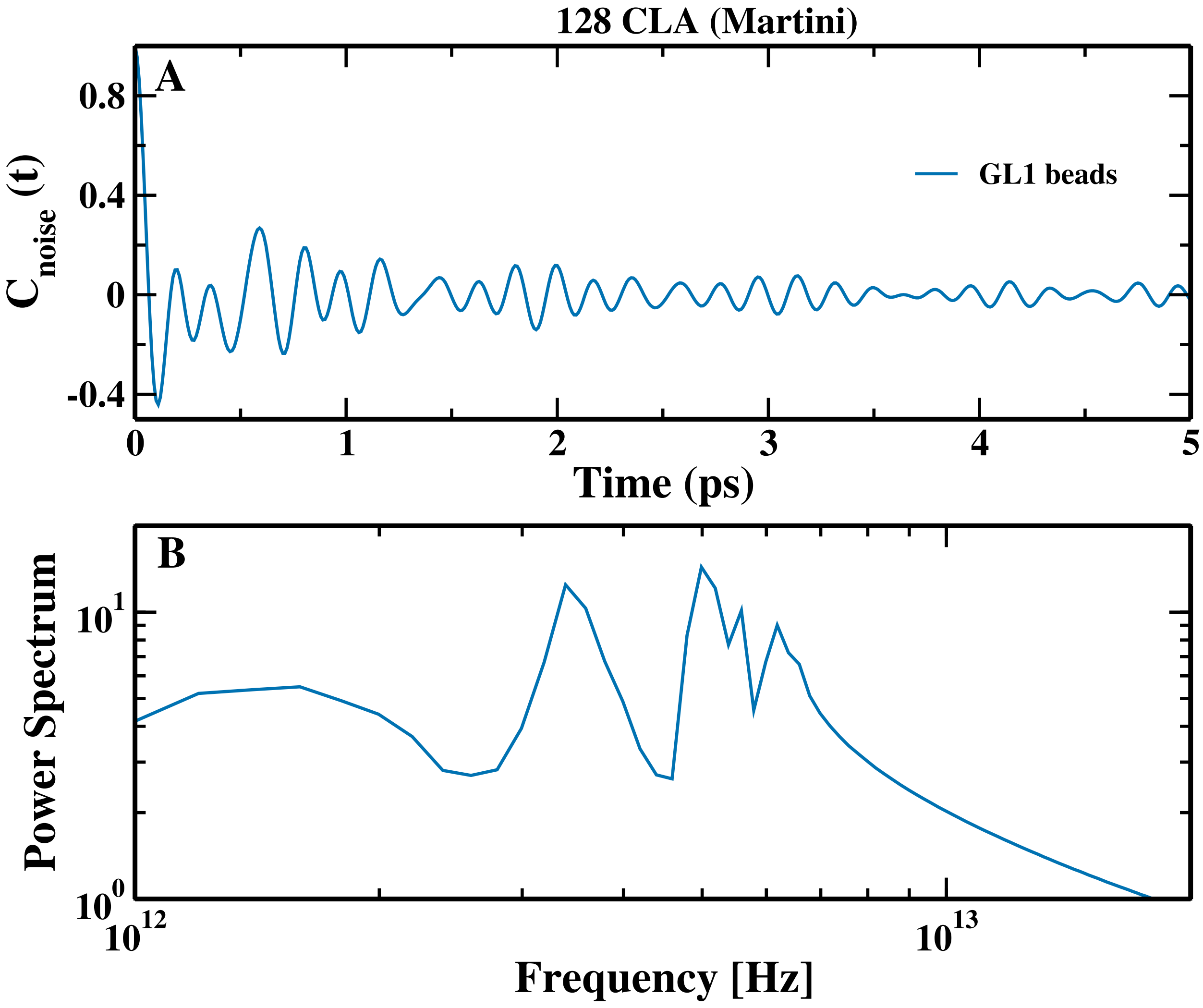


Figure S7: (A) The autocorrelation function, $C_{noise}(t)$, of thermal noise and (B) the power spectrum of thermal noise of lipid beads, GL1, closest to CLA.
